# Topologically associating domains can arise from stochastic folding of heterogeneous fluidlike chromatin

**DOI:** 10.1101/2023.09.12.557077

**Authors:** Luming Meng, Fu Kit Sheong, Qiong Luo

## Abstract

Topologically associating domains (TADs) are critical for gene regulation. Current views attribute TAD formation to cohesin-mediated extrusion and ignore the role of physical properties of *in vivo* chromatin. Here, we demonstrate that the two universal properties: chromatin fluidlike behavior and heterogeneity in DNA-packing density along chromatin, can drive TAD formation. We use DNA-accessibility data to parameterize DNA-packing density along chromatin and simulate stochastic folding of the heterogeneous chromatin in nucleus to yield a conformation ensemble. Such an ensemble can be cross-validated by Hi-C and FISH data. Furthermore, the stochastic folding model allows *de novo* prediction of the establishment and disappearance of key TADs during early T cell differentiation. Together, our work demonstrates that the intrinsic stochastic folding of fluidlike chromatin leads to the prevalence of TAD-like domains in single cells and their cell-to-cell variation, while the heterogeneity in DNA-packing density along chromatin mediates the emergence of TADs at ensemble-averaged level.

**In brief:** A study based on polymer simulation reveals that the two universal physical properties of *in vivo* chromatin fiber: chromatin fluidlike behavior and heterogeneity in DNA-packing density along chromatin play a vital role in TAD formation.

**Highlights:**

- Intrinsic stochastic folding of fluidlike chromatin in nuclear space underlies the prevalence of TAD-like domains in single cells and their cell-to-cell variation
- Heterogeneity in DNA-packing density along chromatin causes the emergence of TADs at ensemble-averaged level
- The disappearance and establishment of key TADs during early T cell differentiation can occur through a stochastic folding process alone, without the need of any cohesin-mediated chromatin extrusion
- The stochastic folding model applies to diverse cell types and is thus able to *de novo* predict the dynamics of genome organization over time

## INTRODUCTION

The physical principles that shape the spatial chromatin organization at the kilobase-to-megabase scale attract much attention(Dekker et al., 2017; Misteli, 2020; Parmar et al., 2019; Rowley and Corces, 2016, 2018). Because such scale covers the typical distances separating enhancers and their distal target promoters, the chromatin organization at such scale may guide the physical interactions between regulatory elements and thus is crucial for gene regulation(Batut et al., 2022; Chen et al., 2018; Dekker and Mirny, 2016; Furlong and Levine, 2018). Through high-throughput chromosome conformation capture (Hi-C) experiments performed on populations of millions of cells, topologically associating domains (TADs) are revealed as a ubiquitous unit of the chromatin organization at such scale in interphase cells (Dixon et al., 2012; Krietenstein et al., 2020; Lieberman-Aiden et al., 2009). Hi-C is a biochemical method to capture long-range chromatin-chromatin contacts (Kempfer and Pombo, 2020; Lieberman-Aiden et al., 2009). The population-averaged Hi-C contact frequency maps show that: (i) TADs are manifested as blockwise patterns on the principal diagonal that represent consecutive self-interacting genomic regions(Lieberman-Aiden et al., 2009; Misteli, 2020; Rowley and Corces, 2018) (Figure 1C); (ii) TADs have sharp boundaries often enriched at cohesin and CTCF binding sites(Dixon et al., 2012; Lieberman-Aiden et al., 2009; Misteli, 2020; Rowley and Corces, 2018) (Figure 1C); and (iii) TADs are relatively conserved across species and cell types(McArthur and Capra, 2021).

**Figure 1.**
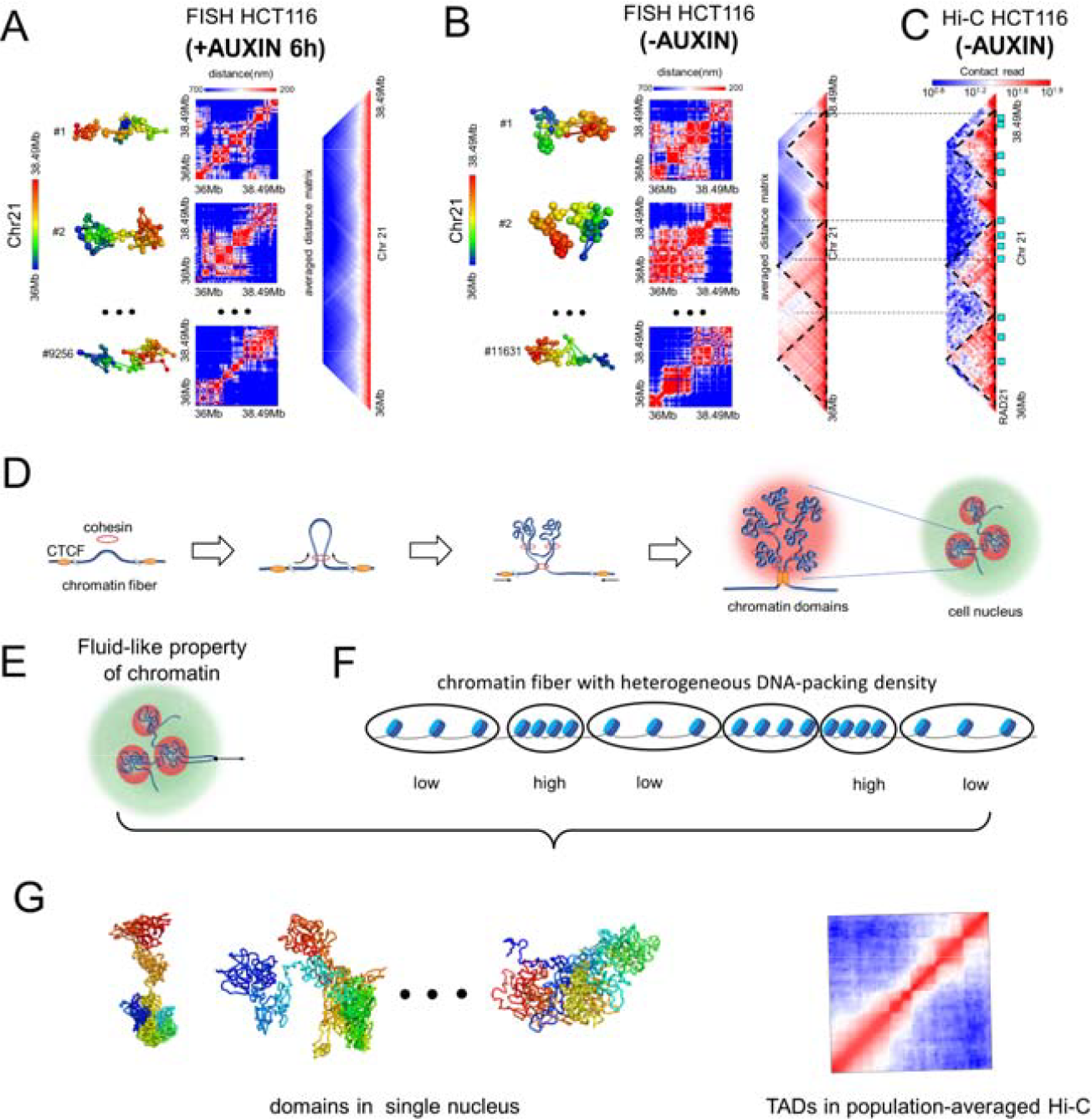
Conflicting evidences in TAD formation and our proposed solution. (A and B) Results of FISH imaging experiments for the 2.5-Mb genomic region (Chr21:36.00 Mb-38.49 Mb) in the transgenic HCT116 cell line with (A) or without (B) auxin treatment which induces cohesin degradation. Left: 3D conformations of the consecutive 30-kb chromatin segments. Middle: Single-cell spatial-distance matrices of 30-kb resolution, showing the widespread presence of domains in individual cells. Right: Averaged spatial-distance matrix. There is no discernable TAD in the averaged spatial-distance matrix of panel (A), while TADs appear as diagonal blocks (indicated by black dashed triangles) in the averaged spatial-distance matrix of panel (B). (C) Population-averaged Hi-C contact map for the same 2.5-Mb genomic region binned at 30-kb resolution. TADs are indicated as black dashed triangles. Cyan squares indicate the positions bound by cohesin (represented by RAD21). A clear correspondence between TAD boundaries and cohesin positions could be observed. (D) Loop extrusion model. Cohesin (red) extrudes chromatin fiber until blocked by CTCF (orange). Several cohesin complexes and a pair of CTCF cooperatively constrain the extruded region as a cross-linked domain/TAD (shaded in red). (E) Recent *in vivo* experiment reveals fluidlike behavior of chromatin(Keizer et al., 2022). (F) *In vivo* chromatin is known to be a heterogeneously condensed DNA polymer(Grigoryev et al., 2009; Ou et al., 2017; Ricci et al., 2015). (G) Proposed stochastic folding model based on the *in vivo* properties shown in (E) and (F). The cell-to-cell variations of chromatin domains (schematically shown on the left) are the consequence of intrinsic random folding of fluidlike chromatin fiber within crowded nuclei. The heterogeneity in DNA-packing density along chromatin leads to the emergence of TADs in the population-averaged Hi-C maps (schematically shown on the right). Calculation details for constructing matrices and identifying TADs boundaries in (A) to (C) can be seen in STAR Methods.

These features of TADs provide a foundation for loop extrusion (LE)(Fudenberg et al., 2017; Fudenberg et al., 2016), which is accepted as the leading principle governing TAD formation(Misteli, 2020; Parmar et al., 2019; Rowley and Corces, 2016, 2018). According to the LE model, a cohesin ring first lands on chromatin and then extrudes chromatin fiber until it encounters obstacles such as CTCF(Misteli, 2020). Meanwhile, other cohesin rings can land on the extruded chromatin fiber to promote chromatin-chromatin interactions(Misteli, 2020) (Figure 1D). Consequently, the extruded chromatin fiber is highly cross-linked into a self-interacting domain; such cross-links limit the degree of freedom of the loci in the domain. By using several *ad hoc* parameters (such as the average residence time of cohesin on chromatin and the cohesin extrusion velocity), LE simulations successfully recapitulate many observations from population-averaged Hi-C maps(Fudenberg et al., 2016; Guo et al., 2022; Nuebler et al., 2018; Vian et al., 2018). However, the cohesin-mediated LE model has failed to account for some observations of chromatin organization at single-cell level(Bintu et al., 2018).

The features of chromatin organization described at single-cell level are quite distinct from TADs(Bintu et al., 2018; Meng et al., 2021; Nagano et al., 2013; Nozaki et al., 2017; Stevens et al., 2017). Although fluorescence in situ hybridization (FISH) imaging experiments have confirmed that interphase chromosomes are indeed partitioned into TAD-like domains, such domains are not conserved across individual cells(Bintu et al., 2018) (Figure 1B). The genomic positions of their boundaries show substantial cell-to-cell variation(Bintu et al., 2018; Meng et al., 2021; Nagano et al., 2013; Nozaki et al., 2017; Stevens et al., 2017) (Figure 1B), and show a preference to reside at CTCF binding sites(Bintu et al., 2018). Such preference leads to the appearance of TADs in the population-averaged Hi-C maps and the enrichment of TAD boundaries at CTCF binding sites. These observations demonstrate that cell-type-invariant TADs are not physical structures that can be seen in single cells, but merely represent an emergent property from cell population averaging. More surprisingly, cohesin depletion, a treatment that abolishes TADs in the population-averaged Hi-C maps, does not alter the widespread presence of TAD-like domains in individual cells(Bintu et al., 2018) (Figure 1A). Furthermore, in the absence of cohesin, domains are reestablished after mitosis(Bintu et al., 2018). These observations pose questions upon the claimed role of cohesin in domain formation and call for re-examination of the LE model (Parmar et al., 2019).

Recent *in vivo* study(Keizer et al., 2022) on the material property of interphase chromatin (e.g. liquid, solid, or gel-like) casts further doubts on the LE model. This study found that subpiconewton force is sufficient to allow a genomic locus to undergo a displacement by several micrometers that is orders of magnitude larger than sizes of TADs (Figure 1E). This result demonstrates that interphase chromatin fibers behave like a free fluidlike polymer rather than a cross-linked polymer gel within the crowded nucleus. The incredible fluidity of *in vivo* chromatin fiber challenges the LE scenario, where the movement of the loci in a self-interacting domain is restricted by the cohesin-mediated cross-links(Misteli, 2020) (Figure 1D). There is a strong need for an alternative model of chromatin domain formation, which should take the material properties of chromatin fiber into account.

Here, we aim to build a novel model for the formation of chromatin domains to reconcile the features of single-cell domains and population-averaged TADs (Figure 1G). This model is based on two universal physical properties of *in vivo* chromatin fiber: fluidlike behavior of the chromatin polymer chain and DNA-packing density heterogeneity along the chain (Figure 1G). We propose that the inherent stochastic folding of fluidlike chromatin within a crowded nucleus results in the prevalence of domains in individual cells. Quite evidently, this stochastic folding process well explains the extreme cell-to-cell variation in the genomic positions of single-cell domain boundaries; but it runs into difficulty in explaining the preference of these boundaries to reside at CTCF binding sites, that is, the key for the emergence of TADs at population level. To solve the problem, we further propose that the heterogeneity of DNA-packing density along *in vivo* chromatin mediates such preference during the stochastic folding process. We propose this hypothesis based on the well-known fact that the structural environment of chromatin is often classified into euchromatin and heterochromatin, which have vastly different DNA-packing density and activity(Babu and Verma, 1987; Bajpai and Padinhateeri, 2020; Grigoryev et al., 2009; Ou et al., 2017; Ricci et al., 2015; Sullivan and Karpen, 2004) (Figure 1F). For the ease of discussion, the term “heterogeneous” chromatin in this paper specifically refers to the chromatin heterogeneous in terms of DNA-packing density.

Together, we build a stochastic folding model, which propose that the inherent stochastic folding of heterogeneous fluidlike chromatin fibers within a crowded nucleus underlies the single-cell domain formation and the emergence of population-averaged TADs. By only using DNA-accessibility data to parameterize the DNA-packing density along the chromatin fiber, we model a heterogeneous chromatin fiber as a heteropolymer and simulate random folding of the heteropolymer under a confined space. Our stochastic folding model excludes any directed process or interaction, including cohesin-mediated chromatin extrusion, attraction of homotypic chromatin to each other (i.e. phase separation), and attraction of heterochromatin to the nuclear lamina. We find that the stochastic folding model can simultaneously recapitulate the features of both the single-cell domains and the population-averaged TADs. Furthermore, the stochastic folding model can quantitatively reproduce the genomic positions of the boundaries of TADs seen in ensemble-averaged Hi-C contact maps and FISH spatial-distance matrices. More importantly, the stochastic folding model allows, for the first time, *de novo* prediction of the dynamic changes in chromatin organization during cell differentiation, by using chromatin accessibility data as the only input. These results reveal that it is the two universal physical properties of 1D chromatin fibers: chromatin fluidlike behavior and heterogeneity in DNA-packing density along chromatin, that play a vital role in the 3D genome organization at the kilobase-to-megabase scale.

## RESULTS

### Description of the stochastic folding model

We build the stochastic folding model to test our hypothesis that inherent stochastic folding of heterogeneous fluidlike chromatin results in the single-cell domain formation and the emergence of population-averaged TADs. We first model the heterogeneous chromatin fiber with a self-avoiding polymer chain consisting of contiguous, coarse-grained beads (Figure 2). Notably, these beads are the same in terms of physical volume but different in terms of their represented DNA sequence lengths, because DNA is heterogeneously condensed into chromatin *in vivo*. The central problem of this formulation lies on how to quantify the represented DNA sequence length (i.e. the DNA-packing density) of each equal-volume bead to develop a heteropolymer model for *in vivo* chromatin fiber. To solve the problem, we turn it into another problem of determining the number *k* of equal-volume beads used to represent a chromatin segment of fixed sequence length. FISH experiments have suggested that there is a clear correlation among DNA-packing density, gene activity, and accessibility of local chromatin segments (Boettiger et al., 2016), namely chromatin segments exhibiting low DNA-packing density always show high activity and high accessibility signals and vice versa. Because DNA accessibility is experimentally measurable through sequencing techniques like DNase-seq or ATAC-seq (Bell et al., 2011; Klemm et al., 2019), we use it as a proxy of the DNA-packing density of a local chromatin segment to determine the value of *k* of the segment (Figure 2 and STAR Methods).

**Figure 2.**
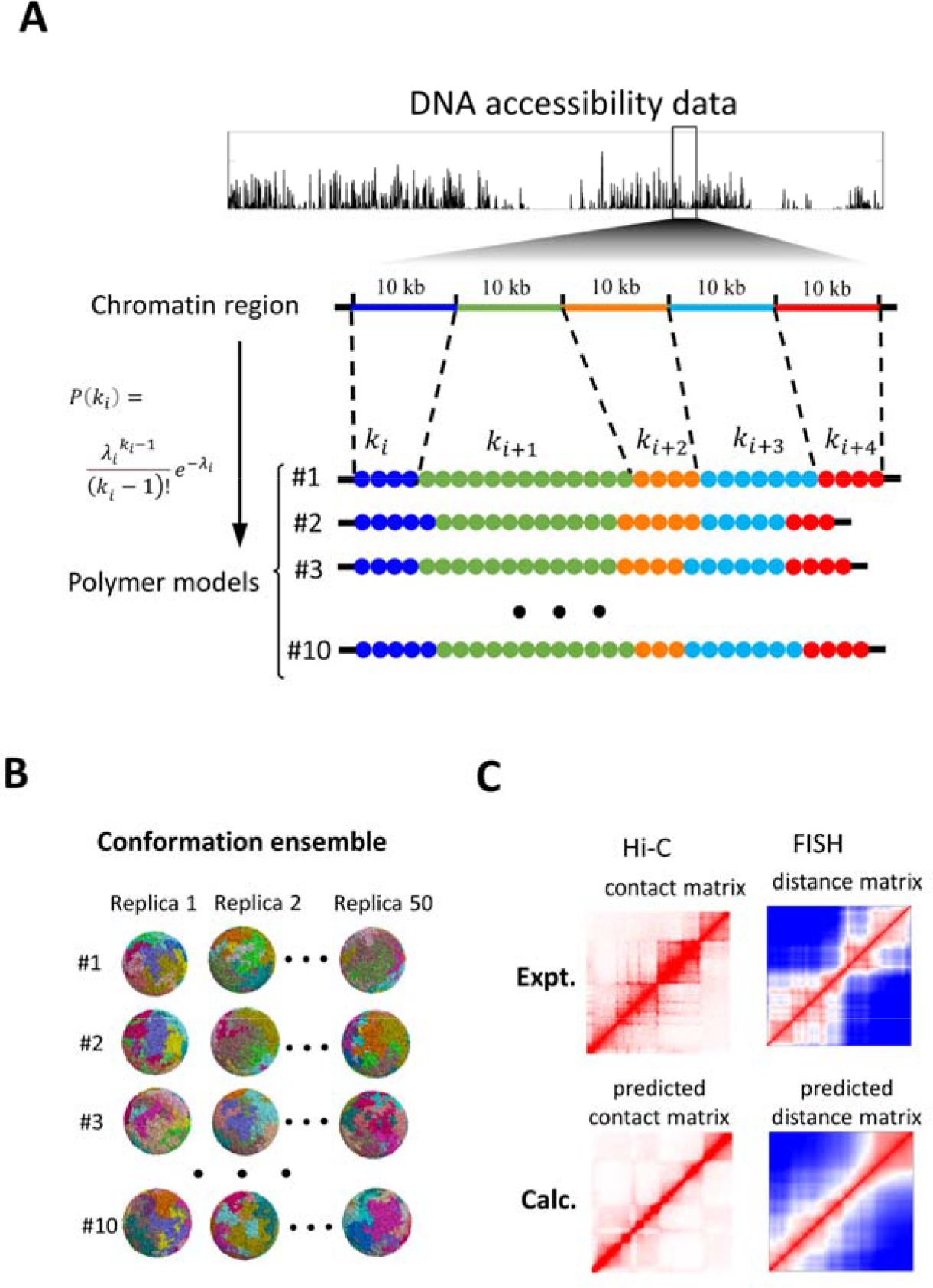
*De novo* prediction of a chromatin 3D conformation ensemble based on chromatin accessibility data. (A) Constructing heteropolymer models for the chromatin fiber of interest. The fiber is partitioned into consecutive 10-kb segments, and the *i*^*th*^ segment is represented with *k*_*i*_ equal-volume beads. A key insight of our approach is that the number *k*_*i*_ is not a constant but is instead modeled as an independent Poisson random variable that is associated with chromatin accessibility of the *i*^*th*^ segment. *P*(*k*_*i*_) is the probability mass function of *k*_*i*_, and λ_*i*_ is the normalized value of DNA accessibility signal of the *i*^*th*^ 10-kb segment. Ten sets of parameters (*k*_*1*_, *k*_*2*_, …*k*_*i*_, *k*_*i+1*_, …) are randomly generated to produce ten heteropolymer models of the same region to model the cell-to-cell variation in DNA-packing density distributions along the chromatin fiber of interest. (B) Results from simulated folding of the 10 heteropolymer models (derived from (A)) in a confined space from 50 different random initial structures which serves as a conformation ensemble for the chromatin region of interest. (C) The conformation ensemble (shown in (B)) can be validated by comparing the predicted contact maps and spatial-distance maps with those experimentally determined by Hi-C and FISH experiments, respectively. Details in the determination of λ_*i*_, drawing *k*_*i*_ from *P*(*k*_*i*_), computational simulation, as well as the construction of contact and spatial-distance matrices from the simulated ensemble can be seen in STAR Methods.

The work flow of establishing the heteropolymer model of *in vivo* chromatin fiber is described in Figure 2. Specifically, we first partition a chromatin region of interest into consecutive 10-kb segments and use *k* equal-volume coarse-grained beads to represent a 10-kb segment (each bead represents (10/*k*) kbs) (Figure 2A). When using experimental population-averaged DNA accessibility data to parameterize the DNA-packing density of each 10-kb segment, we first model *k* as an independent Poisson random variable to account for the observed variation in DNA accessibility among individual cells (Bell et al., 2011; Klemm et al., 2019) (Figure 2A). Next, we randomly generate a group of distinct heteropolymer models for the same chromatin region of interest with varied *k* values for each 10-kb segment (Figure 2 and STAR Methods). These distinct heteropolymer models represent the cell-to-cell variation in DNA-packing density distributions along the chromatin region of interest. For each heteropolymer model, computational simulation is carried out to replicate its stochastic folding process in the confined space where the chromatin volume density is set to 35%(Falk et al., 2019; Ou et al., 2017); such computational simulation is performed for 50 times, each starting from a random initial structure (Figure 2B and STAR Methods). Finally, we gather a conformation ensemble for the modeled chromatin region to compare with the experimental data. In principle, conformations in the ensemble correspond to snapshots of chromatin organization in individual cells and averaging across the conformations in this simulated ensemble yields a population view which can be compared to Hi-C data (Figure 2C). Notably, there are two kinds of sources of experimental population-averaged chromatin accessibility data. One is resulted from ATAC-seq or DNase-seq experiments that are performed on a population of cells. The other is obtained by averaging the accessibility data of single cells provided by single-cell ATAC-seq or DNase-seq experiments. Both kinds of population-averaged chromatin accessibility data would serve as effective input data for our protocol of heteropolymer construction.

Briefly, our stochastic folding model is a polymer-physics-based hypothesis-driven model. This model predicts the conformation ensemble of a chromatin region of interest by using only the population-averaged chromatin accessibility data as input; such conformation ensemble can be compared to not only population-averaged Hi-C data but also single-cell FISH data. Because DNase-seq and ATAC-seq experiments have already yielded a large amount of chromatin accessibility data for diverse cell types (Bell et al., 2011; Klemm et al., 2019), the stochastic folding model enables us to validate our hypothesis of chromatin domain formation in different cell types.

### The stochastic folding model can predict TAD structures at genome-wide scale

We first demonstrate the power of the stochastic folding model by constructing the conformation ensemble of a chromatin region in K562 human erythroleukemia cells. The chromatin region includes 22 autosomes and an X-chromosome. The conformation ensemble for such region is generated using the ATAC-seq data (Dunham et al., 2012) as the input (STAR Methods). For each conformation, we measure contacts between each pair of 10-kb chromatin segments throughout the entire region, and produce the contact matrix of the individual conformation at 10-kb resolution (STAR Methods). We then average these contact matrices of individual conformations across the simulated ensemble to obtain an ensemble-averaged contact matrix, which can be compared to the population-averaged Hi-C contact map(Rao et al., 2015). For quantitative analysis, we identify the chromatin domain boundaries from three kinds of matrices, namely the individual contact matrices, the simulated average contact matrix, and the experimental Hi-C contact matrix in the same way. Since a “boundary” should represent a genomic position displaying a high degree of spatial separation between upstream and downstream chromatin of the stated position, we calculate a separation score at each genomic position to quantitatively describe the degree of the spatial separation of the position (STAR Methods). We define a position as a boundary when the separation score of the position simultaneously satisfies the two criteria: (i) the boundary position should have higher score than any other positions within the 200 kb regions on either side of the position; (ii) the separation score of the boundary should be higher than the baseline obtained by averaging over the whole investigated region including 23 chromosomes.

The analysis of separation score shows that the stochastic folding process alone suffices to produce the prevalence of domain structures within each individual conformation; in addition, the positions of domain boundaries show a large variation among individual conformations (Figure S4C). In fact, the variation is so large that nearly all genomic positions show a nonzero probability to be seen as a domain boundary in the simulated conformation ensemble (Figure 3B). Despite the widespread nonzero distribution, clear peaks in boundary probability are still observed, and they correspond well to chromatin positions with low packing density (Figure 3B). These results are well consistent with the previous observations from FISH experiments(Bintu et al., 2018).

**Figure 3.**
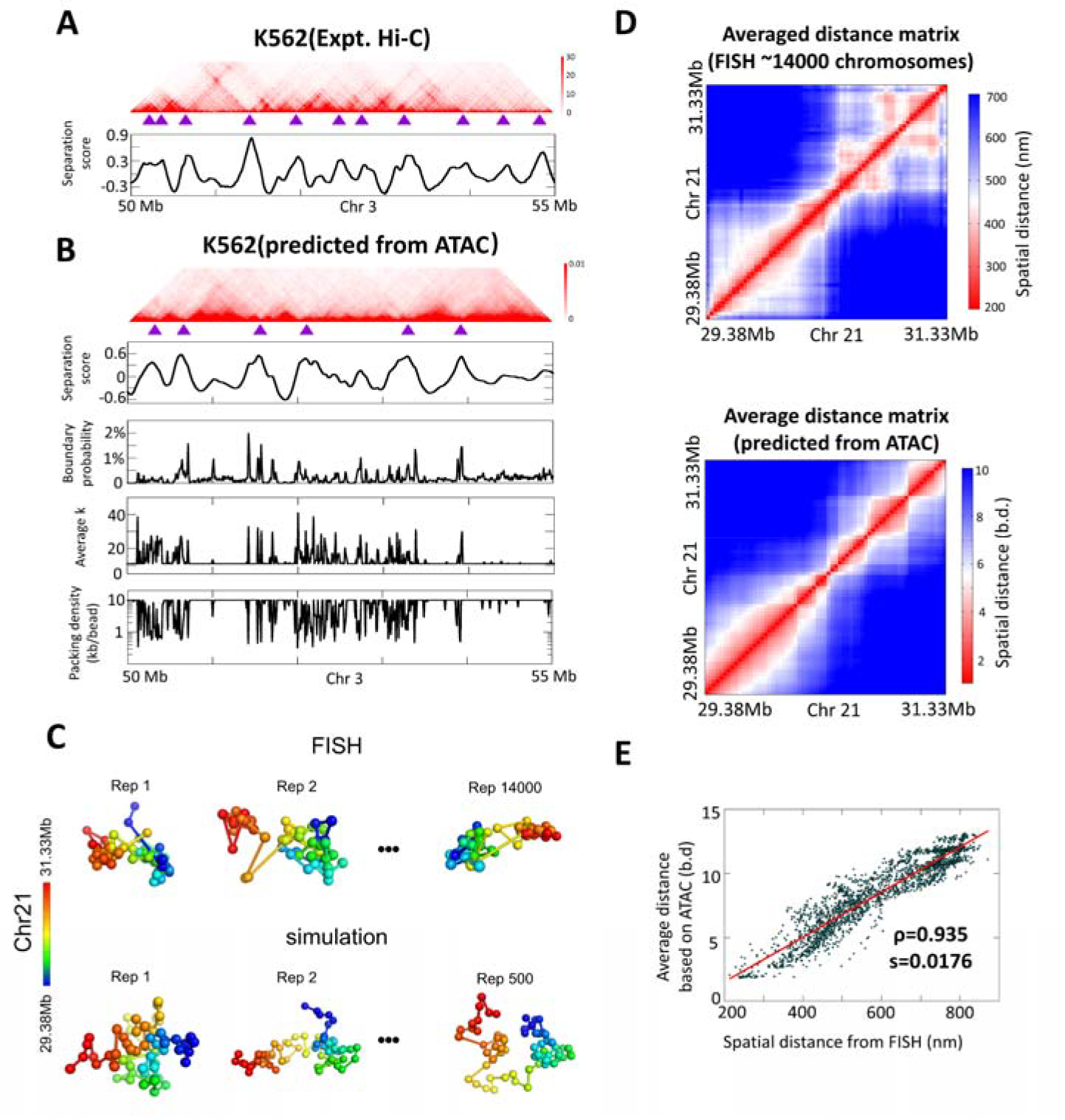
Successful prediction of TAD structures by the stochastic folding model and comparison against experimental Hi-C and FISH data. (A) Top: Population-averaged Hi-C contact matrix for the 5-Mb example region (Chr3:50Mb-55Mb) in the K562 erythroleukemia cell line, binned at 10-kb resolution, with TAD boundaries labeled by purple triangles. Bottom: Separation score for each genomic position. (B) Row 1: Predicted ensemble-averaged contact matrix by the stochastic-folding model using ATAC-seq data as input of the same 5-Mb region shown in (A). TAD boundaries (indicated by purple triangles) are identified by separation score. Row 2: Separation score for each genomic position. Row 3: Probability (fraction of the 500 individual conformations) for each genomic position to appear as a single-cell domain boundary. Row 4: Average *k*_*i*_ for each of the 10-kb segments. Row 5: DNA-packing density of segment along the chromatin, defined by the length of the segment (in kb) divided by the average number of beads in the segment. (C) 3D configurations of the 2-Mb region (Chr21:29.38Mb-31.33Mb) derived from FISH experiment (top) and our simulations (bottom). (D) Top: The experimental averaged spatial-distance matrix at 30-kb resolution across ∼14,000 FISH images of the same 2-Mb region. Bottom: The predicted 30-kb resolution ensemble-averaged spatial-distance matrices of the 2-Mb region. (E) Correlation between the elements of the two spatial-distance matrices shown in panel (D). ρ is the Pearson correlation coefficient and *s* is slope of the fitted line. Details for separation score, construction of various matrices from the simulated ensemble can be seen in STAR Methods. The experimental data is from ref. 20.

In the simulated average contact matrix of the 5-Mb region shown in the example (Chr3:50Mb-55Mb), we see diagonal blockwise patterns with sharp boundaries and the genomic positions of these boundaries align with the TAD boundaries observed in the population-averaged Hi-C map (37) (Figures 3A and 3B). Further comparison throughout the entire region including 22 autosomes and an X-chromosome shows that 1340 (42.2%) out of the total 3176 boundaries predicted by our simulated average contact matrix are seen in the experimental Hi-C map(Rao et al., 2015). However, the number (percentage) decreases sharply to 848 (26.7%) when we randomly select 3176 genomic positions from the entire region instead (Figure S4D). These results demonstrate that the stochastic folding model can quantitatively predict the genomic positions of TAD boundaries at genome-wide scale.

In addition to the validation of the stochastic folding model with Hi-C data, we conduct a further validation using FISH data. We calculate a spatial-distance matrix at 30-kb resolution for each conformation (STAR Methods) and then average them across the ensemble to produce the simulated average spatial-distance matrix. We compare the simulated average spatial-distance matrix of the 2-Mb region (Chr21:29.38Mb-31.33Mb) with the experimental average spatial-distance matrix derived from the published 30-kb resolution FISH images of the same region(Bintu et al., 2018). We find that the two average spatial-distance matrices show a high correlation, with Pearson correlation coefficient of 0.935 (Figures 3D and 3E). Moreover, the conformations of the 2-Mb region in single cells, either provided by the FISH images or by the stochastic folding simulation, unambiguously show the substantial cell-to-cell variation (Figure 3C). These quantitative consistencies provide another support for the validity of the stochastic folding model.

Together, our simulations demonstrate that the conformation ensemble generated by the stochastic folding model can simultaneously recapitulate the population-averaged Hi-C data and single-cell FISH data. Similar TAD boundary predictions have also been successfully made for the chromatin conformation ensembles based on another set of simulations for K562 with DNase-seq data(Dunham et al., 2012) as input (Figures S4 and S6), further illustrating the reliability of our model. Apart from checking the robustness of our stochastic folding model against the use of different experimental input data for parameterization, we have also confirmed the applicability of our procedure on a different cell type, by successfully predict the TAD boundaries for IMR90 lung fibroblasts cell line with ATAC-seq data(Dunham et al., 2012) as input (Figures S5 and S6), again supporting the validity and adequacy of our model.

### The stochastic folding model can predict dynamic changes in chromatin organization during cell differentiation

Previously reported hypothesis-driven models of 3D chromatin organization are heavily focused on the interpretation of results from Hi-C and FISH experiments (Brackey et al., 2020; Fiorillo et al., 2020; Fudenberg et al., 2017; Fudenberg et al., 2016; Yildirim et al., 2022). However, few hypothesis-driven models can trace dynamic changes in 3D genome organization over timescale relevant to mitotic compaction or transcription (which are expected to happen in minutes), let alone biological processes at longer timescales from hours to days (e.g. cell cycle or differentiation), even though understanding these processes from a mechanistic point of view will bring upon a much deeper mechanistic insight into the relationship between genome organization and function(Dekker et al., 2017). The true strength of the stochastic folding model can be better illustrated through a sophisticated example, in which the model correctly predicts the change in chromatin organization during the differentiation of hematopoietic stem and progenitor cells (HSPCs) to CD4 and CD8 double-positive T cells.

This differentiation has attracted considerable attention, owning to the interest in “reprogramming” immune cells for immunotherapies(Li and Turner, 2018). The differentiation process involves a serial of phenotypically well-defined intermediate stages, including multipotent progenitor cells (MPP), common lymphoid progenitor cells (CLP), early T precursor cells (ETP), CD4 and CD8 double negative cells (DN2, DN3, and DN4), and double positive cells (DP). T cell lineage commitment takes place during the DN2-to-DN3 transition(Yui and Rothenberg, 2014). Hu *et al*. systematically investigated the dynamics of chromatin accessibility, chromatin organization, and gene expression of the eight developmental stages, from HSPCs to DP cells(Hu et al., 2018). For each cell type, they profiled the genome-wide average chromatin accessibility data across 1,000 cells by using single-cell DNase sequencing (scDNase-seq); in addition, they measured chromatin-chromatin contacts by performing the Hi-C experiment based on a population of 5,000 to 1,000,000 cells(Hu et al., 2018). They found that *Bcl11b*, encoding a transcription factor critical for T cell commitment, starts to be expressed at the DN2 stage; such an expression is accompanied by an abrupt increase in chromatin interaction frequency in the region containing *Bcl11b*. This increase leads to the appearance of the TAD containing *Bcl11b* in the Hi-C map at the DN2 stage and such *Bcl11b* TAD is further reinforced at the DN4-to-DP transition (Figure 4A). This stepwise and unidirectional change in the chromatin organization of the *Bcl11b*-containing region during differentiation is proposed as an “energy barrier” that locks the fate of a T cell progenitor into the T lineages(Hu et al., 2018). In addition, Hu *et al*. also examined the 2-Mb region (Chr11:18Mb-20Mb) which harbors *Meis1* that encodes a key transcription factor for self-renewal of HSPCs (Figure 4B). They found that there is a delay in the chromatin reorganization of the *Meis1*-contianing region after the T cell lineage commitment, namely the DN2-DN3 transition. The reported Hi-C maps shows that the *Meis1* TAD can be easily identified in the stages from HSPCs to DN3, while a sharp decrease in contact frequency occurs at the DN4 stage and the TAD totally vanishes at the following DP stage(Hu et al., 2018).

**Figure 4.**
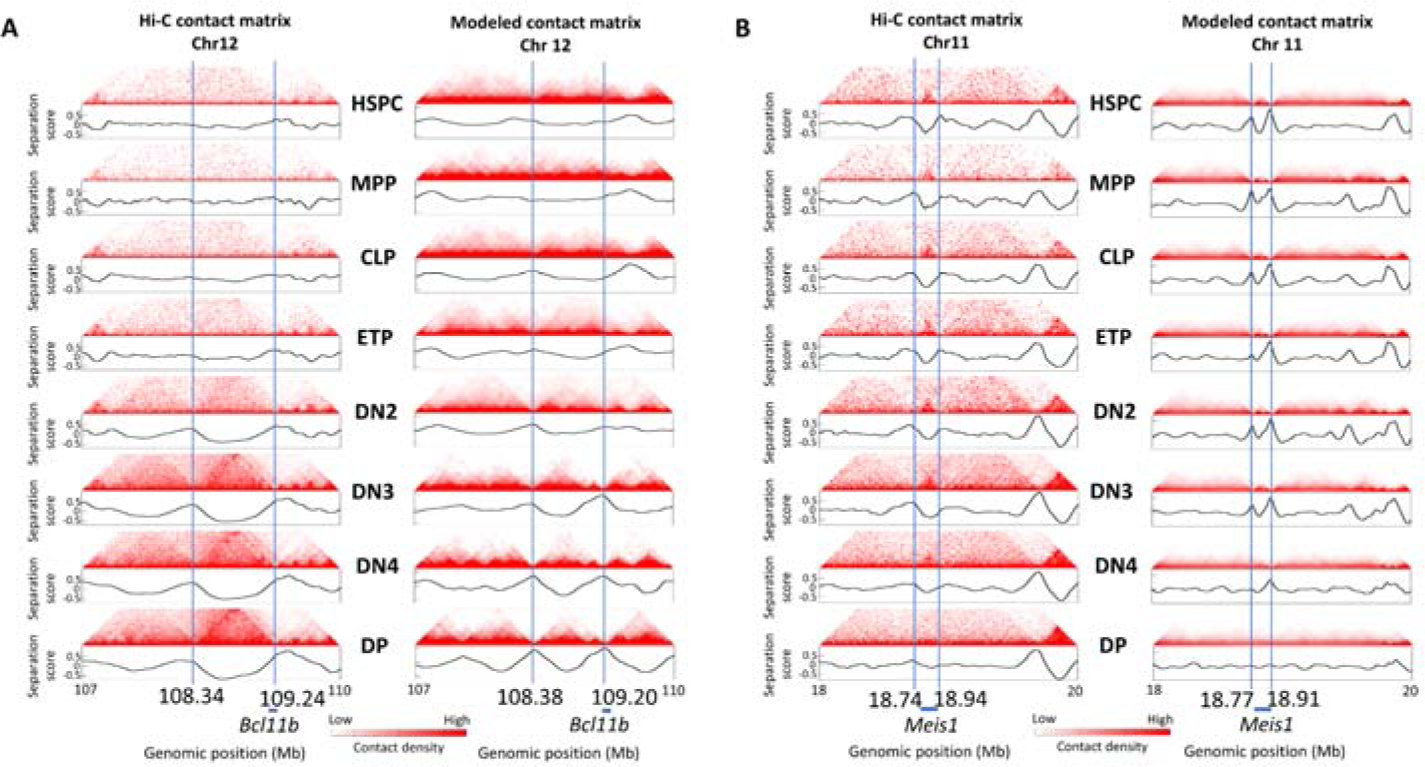
The stochastic folding model allows *de novo* prediction of chromatin reorganization during the differentiation from hematopoietic stem and progenitor cells (HSPCs) to mature immune T cells. (A and B) Left: Hi-C contact-frequency matrices of the two genomic regions containing *Bcl11b* (A) and *Meis1* (B) in cell types of eight developmental stages (HSPC, MPP, CLP, ETP, DN2, DN3, DN4, and DP) during the differentiation (heatmap plot) and separation score for each genomic position (curve plot). Right: The corresponding simulated ensemble-averaged contact matrices of the two same regions (heatmap plot) and separation score for each genomic position (curve plot). All the matrices are of the resolution of 10 kb. Blue lines indicate TAD boundaries identified by separation score. Details for separation score and construction of various matrices from the simulated ensembles can be seen in STAR Methods.

The above-mentioned investigations by Hu *et al*. provide not only the input data (i.e. chromatin accessibility data) for the prediction of conformation ensemble by our stochastic folding model, but also the validation data (i.e. Hi-C data) for the prediction. For each stage, we generate a conformation ensemble by using only the reported average chromatin accessibility data; based on such ensemble, we then construct an ensemble-averaged chromatin-chromatin contact matrix and compare with the published Hi-C contact frequency map (STAR Methods). For quantitative comparison, we again use the separation score to identify TAD boundaries from both the simulated average contact matrices and the published Hi-C maps.

We examine the simulated ensemble-averaged contact matrices of the *Bcl11b*-containing 3-Mb region (Chr12:107Mb-110Mb) from HSPCs to DP cells (Figure 4A). We find that a blockwise pattern containing *Bcl11b* starts to occur at the DN2 stage, with the genomic positions of the two boundaries of the *Bcl11b*-containing block located at 108.38Mb and 109.20Mb, respectively. Notably, the two boundaries of the *Bcl11b*-containing TAD in the Hi-C map separately reside at 108.34Mb and 109.24Mb(Hu et al., 2018). In addition, the values of separation score of the two simulated boundaries increase steadily in the following DN3 to DP stages, indicating a continuous improvement in the strength of the *Bcl11b*-containing block. Comparing the simulated results with the Hi-C maps(Hu et al., 2018), one can immediately see that the stochastic folding model not only precisely reproduces the position and the time of the appearance of the *Bcl11b* TAD, but also correctly predicts the dynamics of the strength of the *Bcl11b* TAD over the cell differentiation process (Figure 4A). We also carry out a similar comparison for the 2-Mb *Meis1*-containing region (Chr11:18Mb-20Mb) (Figure 4B). Again, all changes revealed by the Hi-C maps(Hu et al., 2018) can be precisely reproduced by the predicted conformation ensemble of the stochastic folding model, with the reported chromatin accessibility data as the only input.

All in all, the remarkable success of the stochastic folding simulation in describing the subtle changes of chromatin organization during the differentiation from HSPCs to DP cells is not merely another example that demonstrates the robustness of the stochastic folding model, but actually reveals more intricate details about the mechanism of chromatin organization. More specifically, this example illustrates that the disappearance/establishment of TADs can happen through a stochastic folding process alone, without any aid of cohesin-mediated chromatin extrusion. In addition, the success also implies stochastic folding models built upon different biological states can be compared on the same footing. The comparability across biological states comes from both the folding process we modeled as well as the parameterization procedure. Because the stochastic folding model is only based on the universal physical features of *in vivo* chromatin fiber that holds across different biological states, namely the fluidlike behavior and heterogeneity in DNA-packing density, we do expect the resulting stochastic folding models are universal and comparable. Parameterization-wise, our stochastic folding model uses chromatin accessibility data to parameterize the DNA-packing density along chromatin fiber, and the correlation between DNA-packing density and chromatin accessibility signal of a local chromatin segment is again believed to be conserved across different cell states. Specifically, a chromatin segment exhibiting high accessibility signals is expected to show relatively low DNA-packing density and vice versa. Such universal features and conserved correlation establish the transferability of our model to different cell states, and therefore allows the tracing of dynamic changes in 3D genome organization over time.

### The DNA-packing heterogeneity along chromatin leads to the emergence of TADs in the population-averaged Hi-C maps

Notably, the only parameterization of the stochastic folding model comes from the DNA-packing density along chromatin inferred from DNA accessibility data. We propose that the heterogeneity of DNA-packing density along chromatin is likely to be responsible for the emergence of blockwise patterns (i.e. TADs) in the ensemble-averaged contact maps. To validate the causal relationship between the heterogeneity and blockwise patterns, we further performed simulations to generate conformation ensemble based on a polymer model with each bead representing a constant genomic length, mimicking the case of homogeneous DNA-packing density distribution along chromatin (STAR Methods). While the simulated results show that domain structures could be observed in individual conformations, the probability of each genomic position appearing as a domain boundary across the ensemble is uniformly distributed without significant peaks, and the blockwise pattern is absent in the ensemble-averaged contact matrix (Figure S7). These results suggest that the large variation in domain structures among individual conformations is the outcome of inherent stochastic folding of fluidlike chromatin within a crowded nucleus, no matter whether the fluidlike chromatin is homogeneous or heterogeneous (Figures S4A, S4B, S5A, and S7A). On the other hand, the appearance of TAD structures in ensemble-averaged contact matrix requires heterogeneity in DNA-packing density which would lead to the observed preference of domain boundaries to reside at less densely packed regions (Figures 3B, S4B, S5B)

## DISCUSSION

In this study, we build the stochastic folding model to validate our hypothesis that TADs can arise from intrinsic stochastic folding of heterogeneous fluidlike chromatin within crowded nucleus. We parameterize the DNA-packing density along chromatin by only using chromatin accessibility data and simulate stochastic folding of heterogeneous fluidlike chromatin. Our simulation reveals that the universal physical properties of *in vivo* chromatin fiber, i.e. chromatin fluidlike behavior and heterogeneity in DNA-packing density along chromatin, play a vital role in the chromatin organization at the kilobase-to-megabase scale. The intrinsic stochastic folding of fluidlike *in vivo* chromatin fiber in interphase nucleus causes the prevalence of domains in single cells. The stochasticity of the process shapes the cell-to-cell variation in the genomic positions of domain boundaries. The heterogeneity in DNA-packing density along chromatin mediates preferential residence of domain boundaries at the positions with low DNA-packing density. Such positions correspond to less compacted chromatin regions which are more accessible for proteins such as CTCF. This preference underlies the emergence of TADs in the population-average Hi-C maps.

Our stochastic folding model is in stark contrast with previous hypothesis-driven models of TAD formation, including the LE model(Fudenberg et al., 2017; Fudenberg et al., 2016), the block-copolymer(Fiorillo et al., 2020; Jost et al., 2014) and the Strings&Binders Switch (SBS)(Barbieri et al., 2012; Fiorillo et al., 2020) models. First, these previous models implicitly assume that *in vivo* chromatin is physically constrained and gel-like, by picturing the TAD formation process as *ad hoc* interactions or cross-links between loci. For example, in the LE model, chromatin is highly cross-linked by cohesin because of the assumption that TADs originate from cohesin-mediated chromatin extrusion(Fudenberg et al., 2017; Fudenberg et al., 2016; Misteli, 2020) (Figure 1D). Furthermore, the block-copolymer and the SBS models assume that direct or protein (i.e. binder) mediated interactions between different loci belonging to the same state (i.e. active or inactive) drive the TAD formation(Fiorillo et al., 2020); such homotypic interactions between loci consequently limit their movements. In contrast, our stochastic folding model is established on the experimentally revealed universal feature of *in vivo* chromatin that chromatin is a fluidlike polymer with DNA heterogeneously condensed along it. In the framework of the stochastic folding model, it is the inherent stochastic folding of heterogeneous fluidlike chromatin within crowded nuclei, rather than any *ad hoc* directed extrusion or interaction, that mediates the single-cell domain formation and the emergence of population-averaged TADs. Secondly, the previous models often involve several *ad hoc* input parameters that are currently immeasurable but critical for modeling. For instance, the LE model requires estimations of the average residence time of cohesin on chromatin, the cohesin extrusion speed, and the average separation distance between sites occupied by cohesin(Fudenberg et al., 2017; Fudenberg et al., 2016; Guo et al., 2022). For SBS models, one has to assume types, positions, affinity, and concentration of chromatin binders(Barbieri et al., 2012; Fiorillo et al., 2020). Moreover, it is largely uncertain whether such *ad hoc* parameters show the same values in different cell types(Belokopytova and Fishman, 2021). For example, the PRISMR model, which is one kind of SBS model, infers binder types and positions of the chromatin region containing *EPHA4* locus from the Hi-C data of the region(Bianco et al., 2018). Obviously, it could be problematic to transfer these parameter values to other regions and cell types with potentially different Hi-C patterns. Therefore, these *ad hoc* immeasurable parameters involved in the previous models of TAD formation lead to uncertainty in the transferability of these models to different cell types and hinder the validation of the hypotheses behind these models in different cell types. In contrast, our stochastic folding model does not reply on any *ad hoc* parameter associated with protein-mediated cross-linking/interactions between loci. The only input data of the stochastic folding model is the chromatin accessibility data which is used to parameterize DNA-packing density along chromatin. The chromatin accessibility data can be easily obtained for diverse cell types by ATAC-seq and DNase-seq experiments(Bell et al., 2011; Klemm et al., 2019). Furthermore, high chromatin accessibility signals always mean relatively low DNA-packing density and vice versa, regardless of the cell type under investigation. The robustness of the correlation between chromatin accessibility and DNA-packing density guarantees the transferability of the stochastic folding model and enables the stochastic folding model to practically validate our hypothesis of TAD formation in different cell types. More importantly, the transferability of the stochastic folding model allows the tracing of chromatin organization over time.

Note that the formation of TADs at submegabase scale and A/B compartments at megabase scale are believed to be independent processes in genome organization(Mirny et al., 2019; Nora et al., 2017). The current SF model focuses on TAD formation, and would be unable to reproduce distinct spatial segregation of active euchromatic and inactive heterochromatic regions into A/B compartments. The key contribution of the current model lies on the successful prediction of TAD structures without inclusion of proteins such as cohesin or CTCF, so our model provides a foundation for reinvestigation of the role of proteins (e.g. cohesin) in chromatin organization and shed light to future studies in chromatin conformations.

To conclude, our work demonstrates that inherent stochastic folding of heterogeneous fluidlikechromatin within crowded nuclei can drives the formation of cell-to-cell varying domains in single cells and the emergence of TADs at population level, without any aid of cohesin-mediated chromatin extrusion and interaction between homotypic loci. By using the easily available chromatin accessibility data as the only input data, our work develops the stochastic folding model which enables *de novo* prediction of chromatin organization in different biological states and therefore can map the dynamics of the genome organization over time. The stochastic folding model uncovers the intrinsic stochasticity involved in the chromatin organization at the kilobase-to-megabase scale; such stochasticity reasonably fit the observed variation in gene expression among individual cells(Sood and Misteli, 2022). Our work provides a novel foundation and a powerful tool for gaining deeper mechanistic insights into genome organization and functions.

## STAR⍰METHODS

Detailed methods are provided in the online version of this paper and include the following:

- KEY RESOURCES TABLE
- RESOURCE AVAILABILITY
  ∘ Lead contact
  ∘ Materials availability
  ∘ Data and code availability
- METHOD DETAILS
  ∘ Developing heteropolymer model for chromatin fiber heterogeneous in DNA-packing density
  ∘ Generating random folded conformations of heteropolymer model
  ∘ Conformation ensembles generated in this paper
  ∘ Determination of contact matrices from conformation ensembles
  ∘ Separation score
  ∘ Determination of the pairs of overlapping boundaries among Hi-C contact matrix and the simulated ensemble-average contact matrices
  ∘ Generation of random TAD boundaries
  ∘ Determination of spatial-distance matrices from conformation ensembles
  ∘ Calculation of spatial-distance matrix from fluorescence in situ hybridization experimental data
  ∘ Identifying domain boundaries from the spatial-distance matrix derived from fluorescence in situ hybridization data
  ∘ Reference genome

## Supporting information

Methods part

## SUPPLEMENTARY INFORMATION

Supplemental information can be found online

## ACKNOWLEDGEMENTS

We thank Prof. Han Chen (South China Normal University) and Miss Minhua Lin (South China Normal University) for help with improving the aesthetic of figures. We also thank Dr. Yi Shi(Shanghai Jiao Tong University) and Prof. Xuepeng Chen(Guangzhou Laboratory) for discussions. This work is supported by China Postdoctoral Science Foundation with award number 2022M721203.

## AUTHOR CONTRIBUTIONS

L. M. and Q. L. conceived the project. L. M. conceived and developed the stochastic folding model and performed simulations. L. M. and Q. L. supervised the work. L. M., F. K. S. and Q. L. wrote the manuscript.

