## Supplementary File for "Topologically associating domains can arise from stochastic folding of heterogeneous fluidlike chromatin"

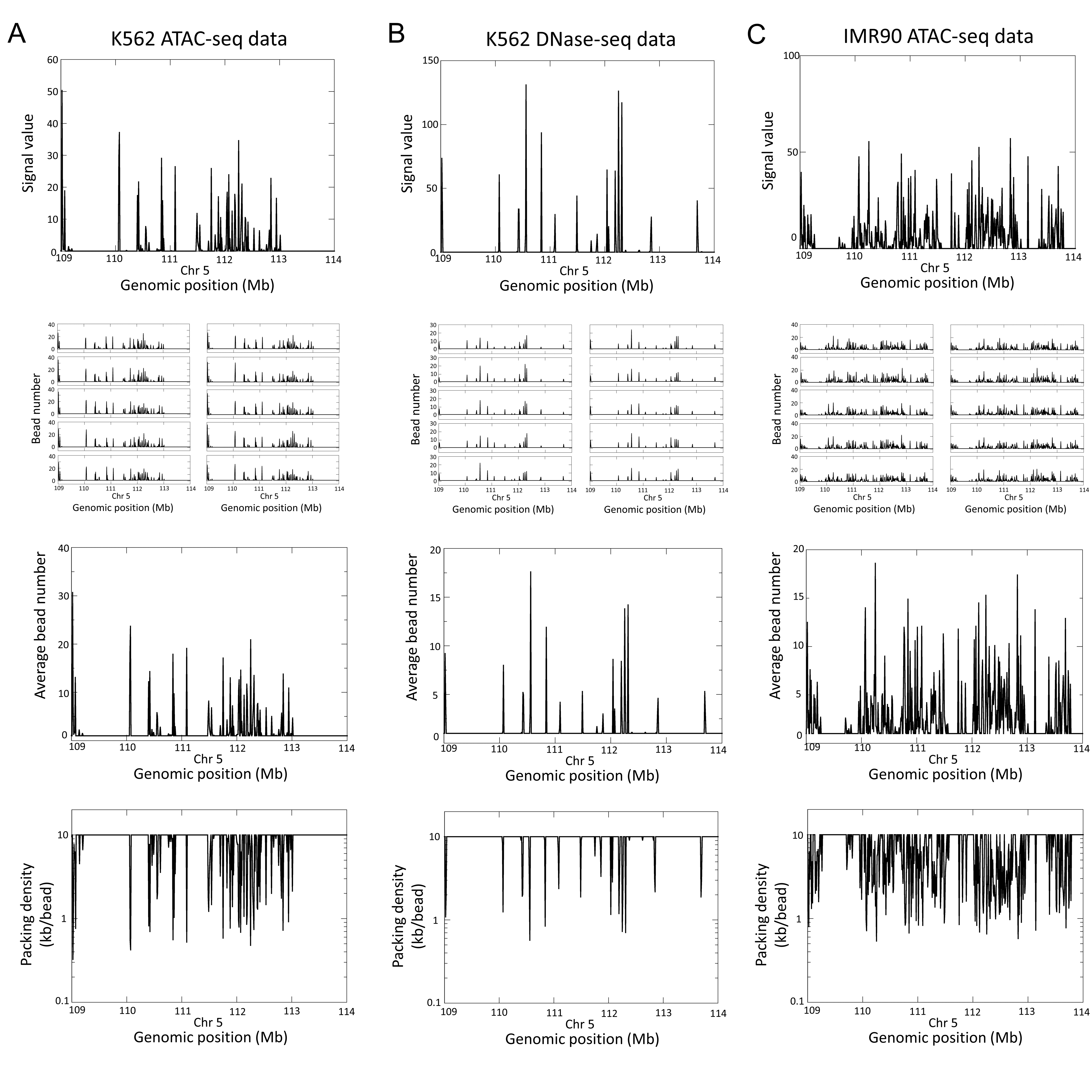

**Figure S1. Plots of bead numbers, different sets of random independent Poisson variables (*k_1_*, *k_2_*,…*k_N_*), and DNA-packing density in our models, resulted from population-based chromatin accessibility data**

(A to C) Row 1: Population-averaged chromatin accessibility data for the 5-Mb chromatin region (Chr5: 109Mb-114 Mb) in K562 erythroleukemia cell type obtained from ATAC-seq experiment (A) or from DNase-seq experiment (B), and in IMR90 lung fibroblast cell type obtained from ATAC-seq experiment (C). Row 2: Ten sets of random independent Poisson variable vectors (*k_1_*, *k_2_*,…*k_N_*) are derived from the data shown in Row 1 by the approach of developing heteropolymer models. Row 3: The mean vector across the ten vectors shown in Row 2, showing the mean value of bead number for each 10-kb segment in the region. Note that the profile of this plot is similar to that of population-averaged chromatin accessibility data shown in Row 1, consistent with our Poisson model of bead number generation. Row4: DNA-packing density, which is defined as the length of the segment (in kb) divided by the average number of beads (shown Row 3).

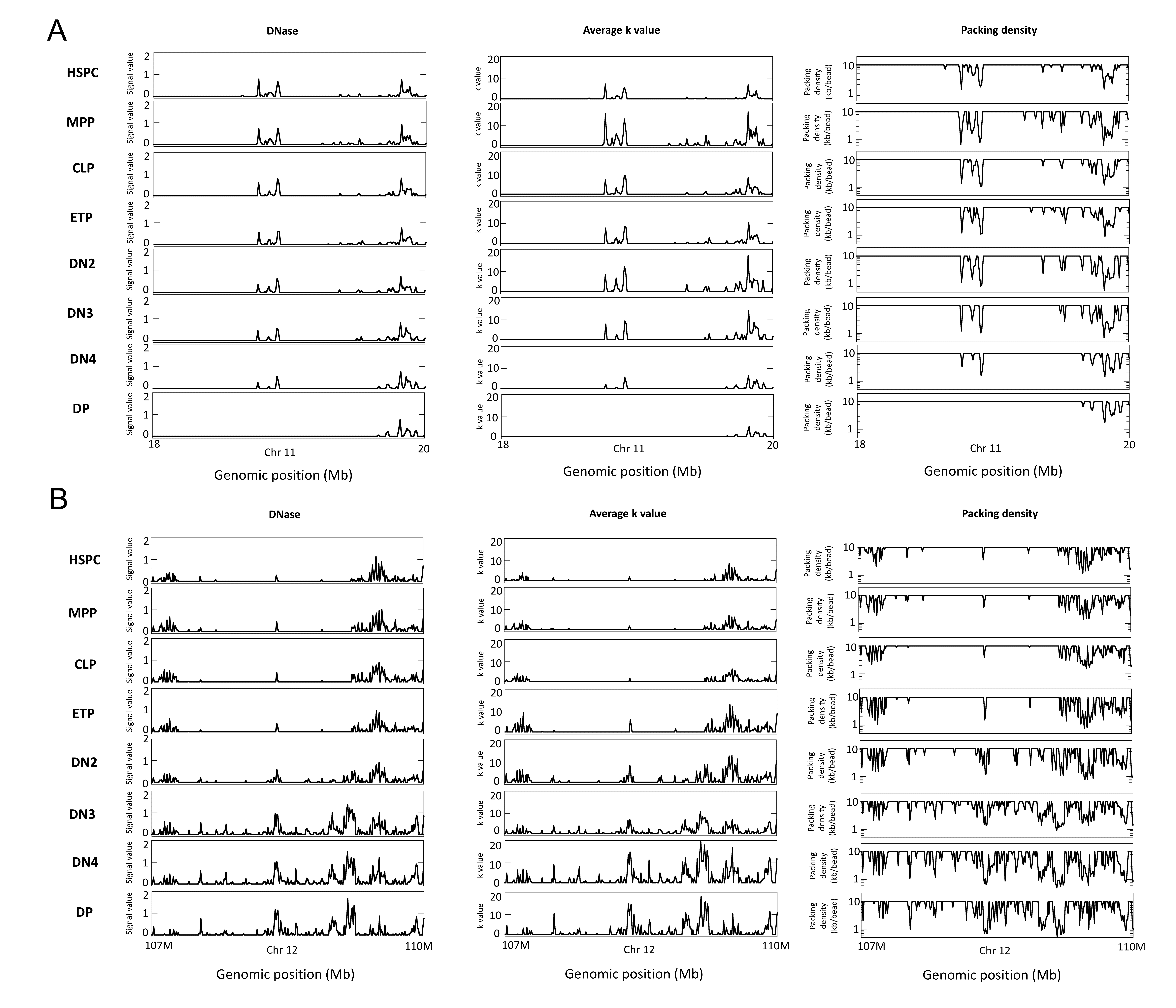

**Figure S2. Plots of population-averaged chromatin accessibility data, averaged bead number distribution, and DNA-packing density for cells at eight developmental stages.**

(A and B) Column 1: Population-averaged chromatin accessibility data for the genomic region (Chr11:18Mb-20Mb) (A) and the region (Chr12:107Mb-110Mb) (B) in cell types of eight developmental stages (HSPC, MPP, CLP, ETP, DN2, DN3, DN4, and DP) during the differentiation shown in Figure 4. Column 2: The mean value of bead number for each 10-kb segment in the region. Column 3: DNA-packing density, which is defined as the length of the segment (in kb) divided by the average number of beads shown in Column 2.

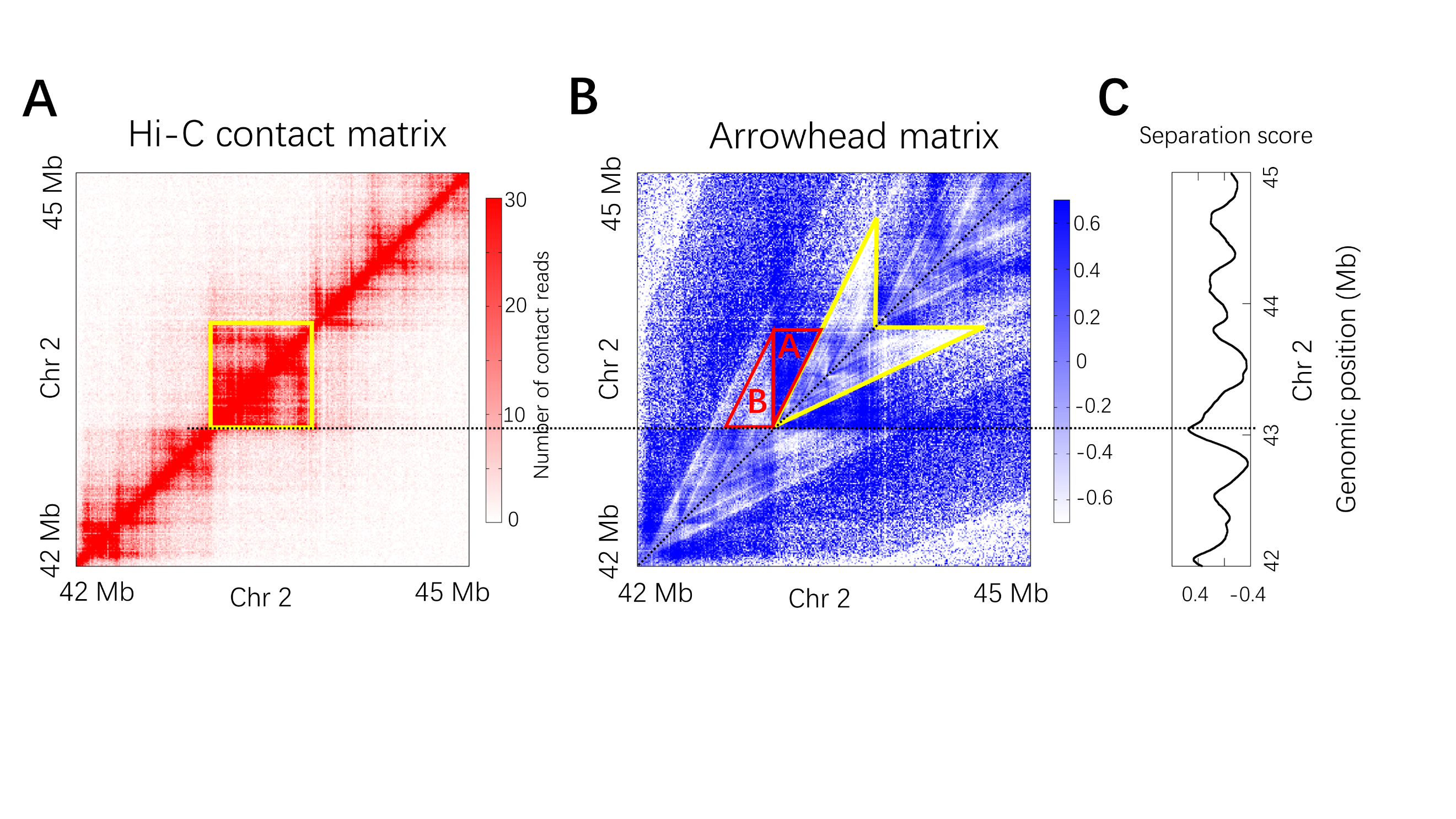

**Figure S3. A scheme of the definition of separation score**

(A)Population-average Hi-C contact frequency matrix for the 3-Mb region (Chr2:42Mb-45Mb) of K562 at 10-kb resolution.

(B)The arrowhead matrix *M* where the arrowhead-shaped motif highlighted by yellow corresponds to the yellow highlighted domain square in panel A.

(C) Separation score of a position along the diagonal of the arrowhead matrix is defined based on the two constructed edge-shared congruent right triangles at that position. As an example, such two edge-shared congruent right triangles of the position corresponding to a domain boundary are highlighted by red in panel B. The length of horizontal edge of the two congruent right triangles is 250-kb and the length of the vertical edge (or shared edge) is 500-kb.

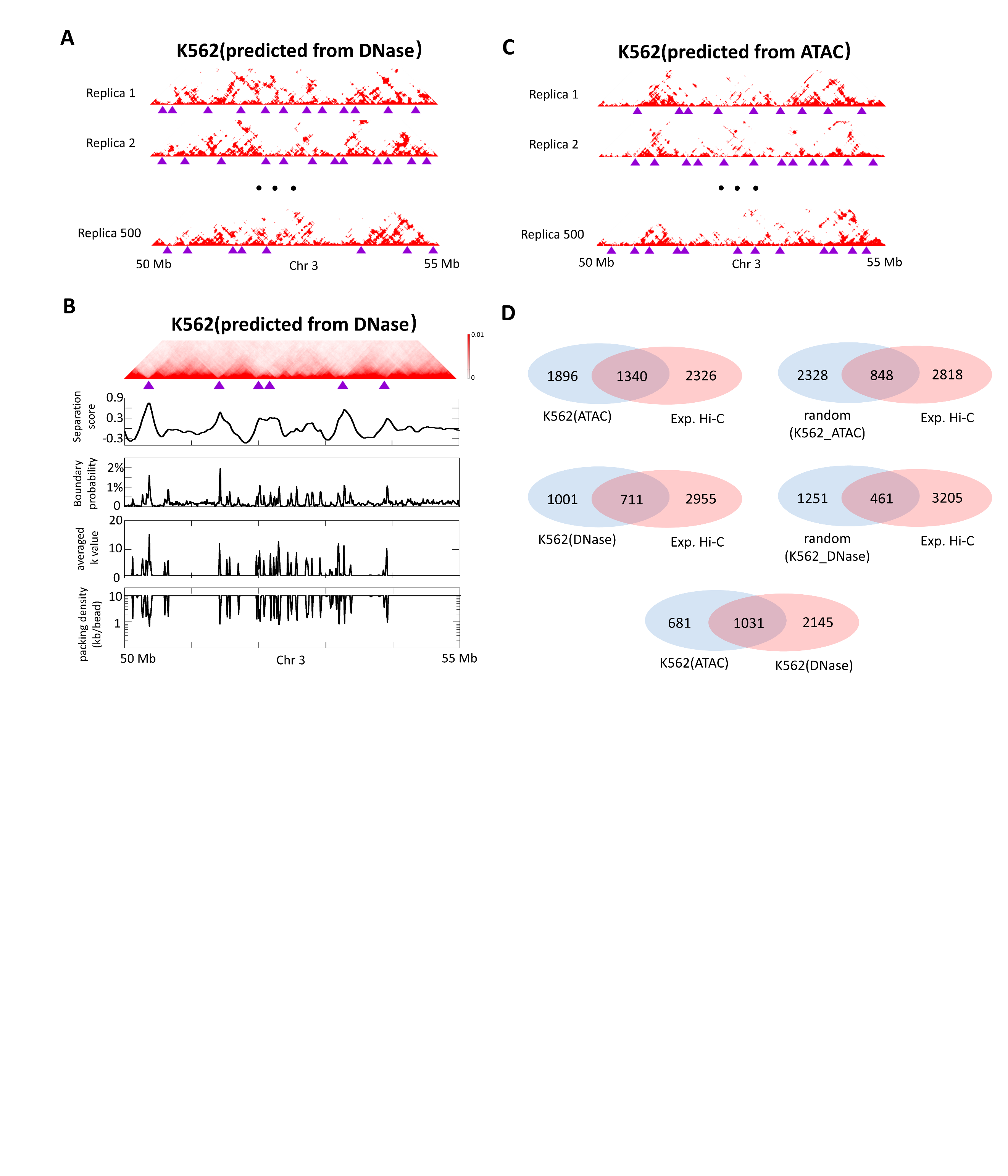

**Figure S4. Prediction of chromatin organization of K562 cell type by the SF model and comparison against experimental Hi-C data**

(A)Individual contact matrices of the 5-Mb chromatin region (Chr3:50Mb-55 Mb) in K562, calculated from individual conformations in the conformation ensemble of K562 derived from DNase-seq data. Domain structures are widespread in individual conformations and their boundaries (indicated by purple triangles) show apparent variation among conformations.

(B)Row 1: Ensemble-averaged contact matrix across the 500 individual contact matrices shown in A. Predicted TAD boundaries are indicated by purple triangles. Row 2: Separation score plot for segments in this chromatin region. Row 3: Probability (fraction of the 500 individual conformations) for each genomic position to appear as a single-cell domain boundary. Row 4: Averaged *k_i_* for each of the 10-kb segments. Row 5: DNA-packing density along the segment, defined by the length of the segment (in kb) divided by the average number of beads in the segment.

(C)Contact matrices of the same 5-Mb chromatin region in K562, calculated from individual conformations in the conformation ensemble of K562 derived from ATAC-seq data. Domain structures are widespread in individual conformations and their boundaries (indicated by purple triangles) show apparent variation among conformations.

(D)Analysis of the overlap of TAD boundaries among Hi-C contact matrix and the two ensemble-averaged contact matrices calculated from the two ensembles of K562, throughout the region including 22 autosomes and X-chromosome of K562.

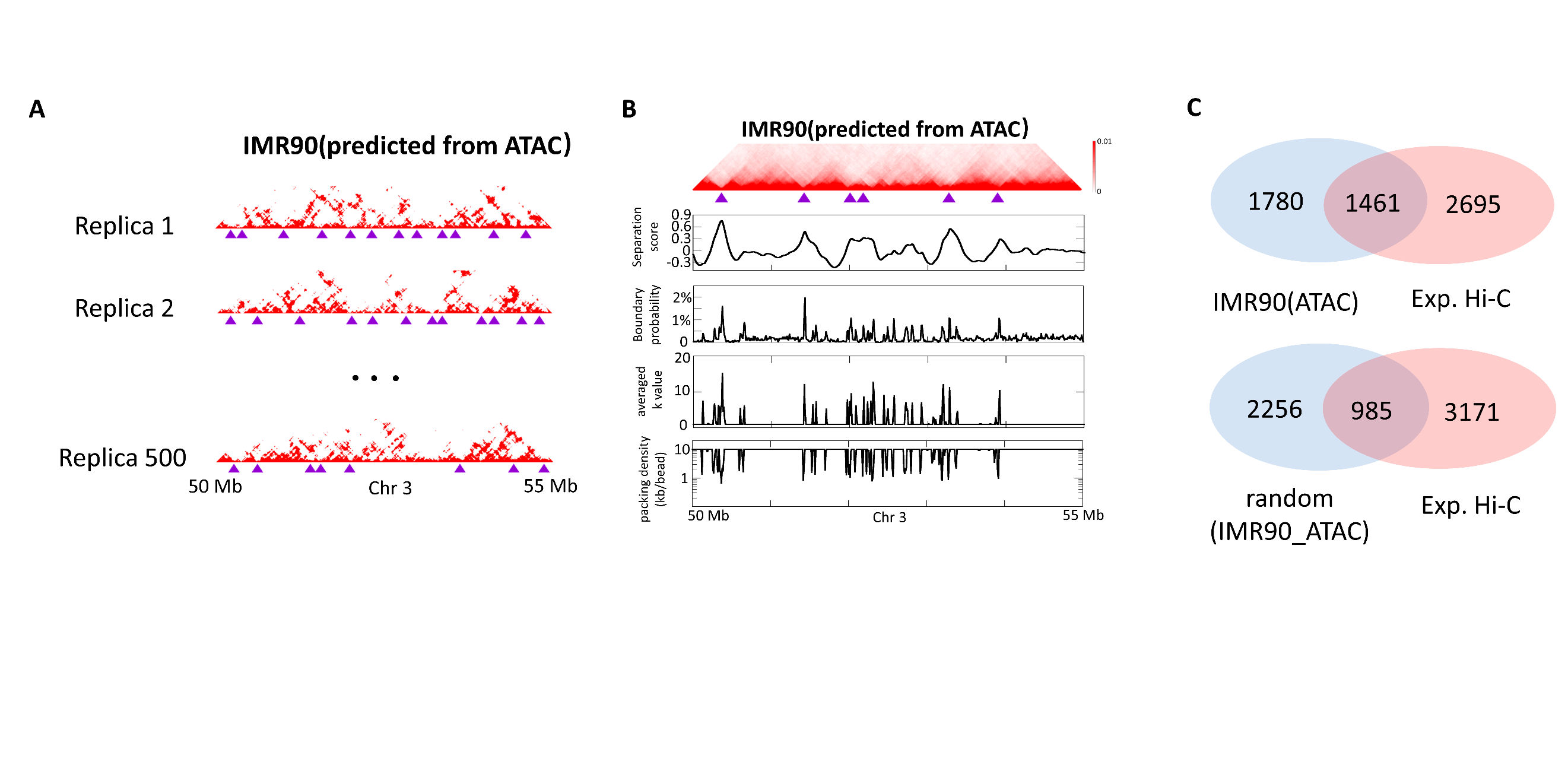

**Figure S5. Prediction of chromatin organization of IMR90 cell type by the SF model and comparison against experimental Hi-C data**

(A)Individual contact matrices of the 5-Mb chromatin region (Chr3:50Mb-55 Mb) in IMR90, calculated from individual conformations in the conformation ensemble of IMR90 derived from ATAC-seq data. Domain structures are widespread in individual conformations and their boundaries (indicated by purple triangles) show apparent variation among conformations.

(B)Row 1: Ensemble-averaged contact matrix across the 500 individual contact matrices shown in A. Predicted TAD boundaries are indicated by purple triangles. Row 2: Separation sore plot for segments in this chromatin region. Row 3: Probability (fraction of the 500 individual conformations) for each genomic position to appear as a single-cell domain boundary. Row 4: Averaged *k_i_* for each of the 10-kb segments. Row 5: DNA-packing density along the segment, defined by the length of the segment (in kb) divided by the average number of beads in the segment.

(C)Analysis of the overlap of TAD boundaries among Hi-C contact matrix and the ensemble-averaged contact matrix calculated from the ensemble of IMR90, throughout the region including 22 autosomes and X-chromosome of IMR90.

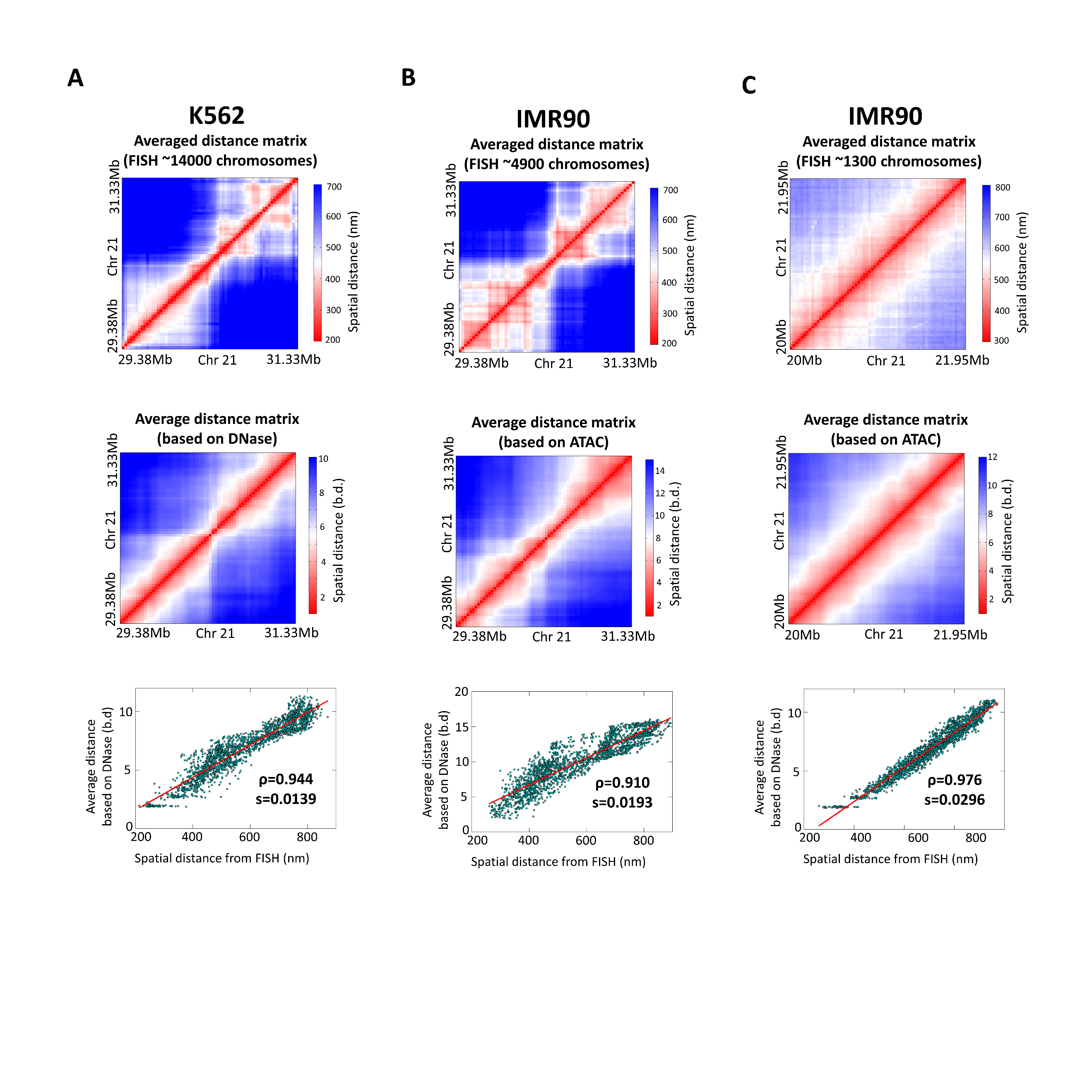

**Figure S6. Predictions of chromatin organization by the SF model and comparison against experimental FISH data**

(A to C) Row 1: The experimental averaged distance matrices at 30-kb resolution across ~14,000 FISH images of the 2-Mb region (Chr21:29.38Mb-31.33Mb) in K562 cell type (a), across ~4900 FISH images of the 2-Mb region (Chr21:29.38Mb-31.33Mb) in IMR90 cell type (b), across~1300 FISH images of the 2-Mb region (20Mb-21.95Mb) in IMR90 cell type (c). Row 2: The 30-kb resolution ensemble-averaged distance matrix of the region shown in Row 1, calculated from the conformation ensemble of K562 predicted from DNase-seq data (a) or calculated from the conformation ensemble of IMR90 predicted from ATAC-seq data (b and c). Row 3: Correlation between the elements of the FISH distance matrix shown in Row 1 and the ensemble-averaged distance matrix shown in Row 2.

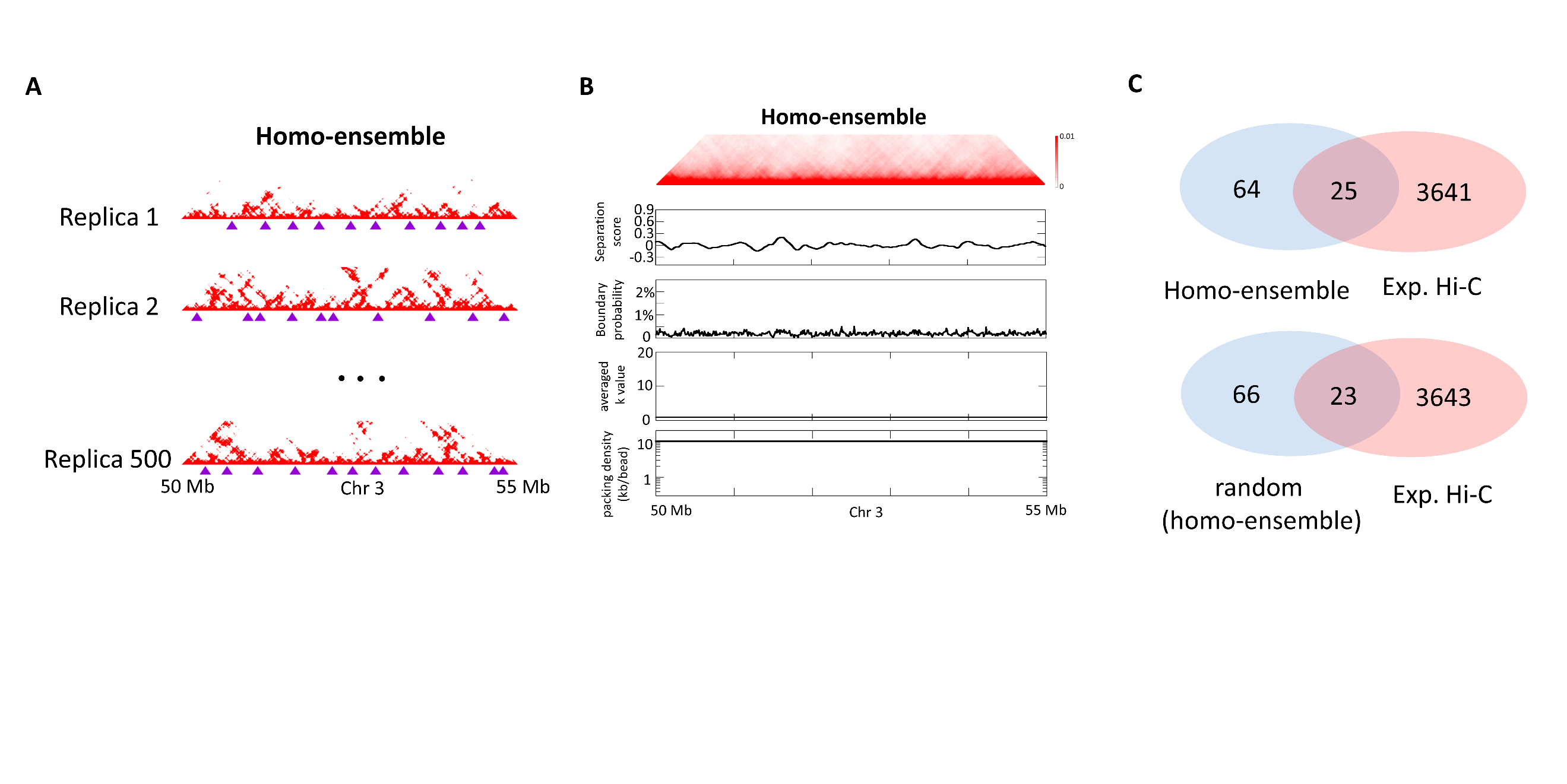

**Figure S7. Domain boundaries in the conformation ensemble generated based on the polymer model that is homogeneous in DNA-packing density**

(A)Individual contact matrices of the 5-Mb chromatin region (Chr3:50Mb-55 Mb), calculated from individual conformations in the conformation ensemble of homogeneous DNA-packing density model. Domain structures are widespread in individual conformations and their boundaries (indicated by purple triangles) show apparent variation among conformations.

(B)Row 1: Ensemble-averaged contact matrix across the 500 individual contact matrices shown in a. No discernable diagonal blocks appear in the 5-Mb region. Row 2: Separation sore for each genomic position. Row 3: Probability (fraction of the 500 individual conformations) for each genomic position to appear as a single-cell domain boundary. Row 4: Averaged *k_i_* for each of the 10-kb segments. Row 5: DNA-packing density along the segment, defined by the length of the segment (in kb) divided by the average number of beads in the segment.

(C)Analysis of the overlap of TAD boundaries among Hi-C contact matrix (of K562 as an example) and the ensemble-averaged contact matrix calculated from conformation ensemble of homogeneous DNA-packing density model.
