## Supplementary material for "Topologically associating domains can arise from stochastic folding of heterogeneous fluidlike chromatin": Methods part

**STAR★METHODS**

**Key resources table**

| REAGENT or RESOURCE | SOURCE | IDENTIFIER |
| --- | --- | --- |
| Deposited data | | |
| Simulated conformation ensembles for K562, IMR90 and eight developmental stages from HPSC cells | This paper | https://ngdc.cncb.ac.cn/bioproject/browse/PRJCA017011 |
| Software and algorithms | | |
| Code for the SF model | This paper | https://github.com/TheMengLab/Stochastic-folding |
| Other | | |
| Population average Hi-C data for IMR90 and K562 |  | GEO:GSE63525 |
| ATAC-seq data for K562 |  | ENCODE:ENCSR483RKN |
| Population average Hi-C data for HCT116 |  | GEO:GSM2795535 |
| RAD21 data for HCT116 |  | GEO:GSM5518543 |
| ATAC-seq data for IMR90 |  | GEO:GSE169767 |
| DNase-seq data for K562 |  | GEO:GSM1008601, GEO:GSM1008580,  GEO:GSM1008567,  GEO:GSM816655,  GEO:GSM1008602,  GEO:GSM1008558 |
| 3D FISH data for K562, IMR90, HCT116(AUXIN treated) and HCT116(AUXIN untreated) | B. Bintu et al. |  |
| Hi-C data and DNase-I hypersensitivity sites data for eight developmental stages from HPSC cells |  | GEO:GSE79422 |

**RESOURCE AVAILABILITY**

**Lead contact**

**Materials availability**

No new materials were generated during the course of this study.

**Data and code availability**

- All original code for the SF model has been deposited on https://github.com/TheMengLab/Stochastic-folding. Simulated conformation ensembles are available on https://ngdc.cncb.ac.cn/bioproject/browse/PRJCA017011
- Any additional information required to reanalyze the data reported in this paper is available from the lead contact upon request.

**METHOD DETAILS**

**Developing heteropolymer model for chromatin fiber heterogeneous in terms of DNA-packing density**

We first partition the chromatin region of interest into *N* consecutive 10-kb length segments. Next, we model the number *k_i_* of contiguous, coarse-grained, and equal-volume beads used to represent the *i^th^* segment as a random independent Poisson variable. The probability of drawing a particular value of *k_i_* from the distribution, *P*(*k_i_*), is defined as:

$P\left( k_{i} \right)=\frac{{\lambda_{i}}^{k_{i}}}{k_{i}!}e^{{-\lambda}_{i}}$	(s1)

The definition of λ_i_. Based on the experimental population-based chromatin accessibility data of the chromatin region of interest (i.e. DNase-seq or ATAC-seq data), we calculate the DNA accessibility signal ζ for each 10-kb segment within the chromatin region of interest. *λ_i_* is the value of DNA accessibility signal of the *i^th^* 10-kb segment normalized by the median of those detected signals, and is defined as:

$\lambda_{i}=\frac{\zeta_{i}}{\text{Median}(\zeta)}$	(s2)

The determination of *k_i_*. With *λ_i_* of certain value, the value of *k_i_* is determined by using a pseudo-random number generating function capable of drawing Poisson random variables with the formula of (Eq. s1). When a particular value of *k_i_* is drawn (say *k_i_* = 2), we model the corresponding segment with *k_i_* +1 beads (say 3 beads). For the clarity of later discussion on the distribution and property of *k_i_* we would leave out the +1 in the text but we hereby assume that to be implicitly implied when we convert *k_i_* to the number of beads in a particular polymer model.

Generating a group of polymer models. Performing the process of determining *λ_i_* and *k_i_* for each 10-kb segment in the chromatin region of interest, we obtain a certain set of (*k_1_*, *k_2_*,…*k_N_*) to build a heteropolymer model with a certain DNA-packing density distribution. To model the cell-to-cell variation in the DNA-packing density distribution, we generate ten distinct sets of (*k_1_*, *k_2_*,…*k_N_*) which represent ten different distributions of DNA-packing density. Namely, we build ten distinct heteropolymer models for the chromatin region of interest. Figure S1 shows the resulted ten sets of (*k_1_*, *k_2_*,…*k_N_*) for the 5-Mb chromatin regions (Chr5:109Mb-114 Mb) of K562 and IMR90 as examples. For simulations that mimics the random folding of homogeneous chromatin fiber, we set all the Poisson variables in (*k_1_*, *k_2_*,…*k_N_*) to be 0. Namely, for homogeneous chromatin fiber, each 10-kb segment is represented by one equal-volume bead.

**Generating random folded conformations of heteropolymer model**

The polymer model is generated via a hierarchical process, starting from a highly coarse-grained model and slowly refine to the targeted 10kb resolution model. A Brownian-like process is used in the generation of a feasible model, in which the velocity of each bead is randomly generated at every integration step following the corresponding Maxwell-Boltzmann distribution.

Numerical integration. Following the mentioned method of developing heteropolymer model, each chromosome is represented as a polymer chain consisting of contiguous, equal-volume beads with the diameter of *b*. The 3D folding of the chromatin region of interest is simulated by numerical integration starting from an initial conformation for certain steps with the size $\Delta t$ for each step. $\Delta t$is defined as:

$\Delta t=\sqrt{0.2b*\frac{m}{F_{max}}}$	(s3)

where $F_{max}$ is the maximum force among the forces applied on beads in the polymer model of the chromatin region of interest. The diameter *b* and the mass *m* of each bead are set to 1 arbitrary unit. The conformation of simulated model is updated with the formula:

$\vec{r}_{i,t+\Delta t}=\vec{r}_{i,t}+\vec{v}_{i,t}\Delta t+\frac{\vec{f}_{i,t}}{2m}{\Delta t}^{2}$	(s4)

where $\vec{r}_{i,t}$ and $\vec{r}_{i,t+\Delta t}$ separately represent the coordinates of the *i^th^* bead before and after the integration step. The velocity vectors $\vec{v}_{i,t}$ are drawn from the Gaussian distribution at every integration step such that the magnitude of the momentum follows from the Maxwell-Boltzmann distribution. $\vec{f}_{i,t}$ represents the force applied on the *i^th^* bead based on the potential energy described below. $m$ is the mass of each bead with the value of 1 and $\Delta t$ is the step size defined in Eq. s3.

Potential energy used in numerical integration. In each step of numerical integration, we define the total potential energy of a conformation with three terms that are described below:

(i) Harmonic oscillator potential energy, $E_{har}\left( r_{n,n+1} \right)$, is introduced to ensure the connectivity of chromosome backbone and is calculated according to the formula as the following:

$E_{har}\left( r_{n,n+1} \right)={\frac{1}{2}k_{b}\left( r_{n,n+1}-b \right)}^{2}$	(s5)

where $r_{n,n+1}$ is the Euclidean distance between pair of the sequentially adjacent *n^th^* and (*n*+1)*^th^* beads. The equilibrium distance between the *n^th^* and (*n*+1)*^th^* beads is equal to the bead diameter *b* with the value of 1 arbitrary unit. The force constant $k_{b}$ is defined as:

$k_{b}=\frac{200kT}{b^{2}}$	(s6)

where $k$ is the Boltzmann constant, *T* the temperature, and *b* the diameter of bead with the value of 1 arbitrary unit. Here, *kT* is considered as a unit amount of energy with value of 1 arbitrary unit.

(ii) Repulsive potential energy, $E_{WCA}\left( r_{n,m} \right)$, is introduced to avoid spatial overlapping between pair of the non-sequentially-adjacent *n^th^* and *m^th^* beads and is calculated according to the following:

$E_{WCA}\left( r_{n,m} \right)=\left\{ \begin{matrix} 4\varepsilon\left( \left( \frac{\sigma}{r_{n,m}} \right)^{12}-\left( \frac{\sigma}{r_{n,m}} \right)^{6} \right), if r_{n,m}\leq2^{1/6}\sigma\\ -\varepsilon, else \end{matrix} \right.$	(s7)

where $r_{n,m}$ represents the Euclidean distance between the *n^th^* and the *m^th^* beads and $\sigma$ and $\varepsilon$ are the two constants with $\sigma=b$ and $\varepsilon=\frac{1}{2}kT$.

(iii) Spherical restraint potential energy,$E_{sphere}\left( r_{n} \right)$, is introduced to constrain the folding of the polymer model of chromatin in a confined spherical space like cell nucleus. During the modeling process, the center of the spherical space is fixed at the origin of coordinates (0,0,0). The radius of the sphere $r_{s}$ is defined as:

$r_{s}=\frac{1}{2}b\left( N/0.35 \right)^{1/3}$	(s8)

where *N* is the total number of beads in the polymer model and *b* the diameter of each bead with the value of 1. The value of 0.35 is adopted to set the value of chromatin volume density at 35%. Based on the definition of the spherical space, spherical restraint potential energy,$E_{sphere}\left( r_{n} \right)$, is calculated according to the formula:

$E_{sphere}\left( r_{n} \right)=\left\{ \begin{matrix} \frac{r_{n}-r_{s}}{b}kT, if r_{n}\geq r_{s} \\ 0, else \end{matrix} \right.$	(s9)

Where $r_{n}$ is the Euclidean distance between the *n^th^* bead and the center of the sphere space (i.e. origin of coordinates).

Hierarchical calculation strategy. As mentioned previously, we equally partition the chromatin region of interest into *N* consecutive 10-kb segments and build a bead-on-a-string representation of the region. We define the resolution of the representation as the sequence length of individual partitioned segments, namely 10 kb. To rapidly achieve the 3D conformation of the chromatin region of interest at 10-kb resolution, we employ a hierarchical protocol to first generate structure of low resolution and then convert the low resolution structure to high resolution one for further optimization. The hierarchical protocol includes the following steps:

(i) Determination of the number of beads used to represent the chromatin region of interest at hierarchical resolutions. We first determine the number *k* of beads used to represent each 10-kb segment according to the above mentioned method of developing heteropolymer model and obtain a certain vector (*k_1_*, *k_2_*,…*k_N_*) to build a bead-on-a-string representation for the chromatin region of interest at 10-kb resolution. Next, we sequentially merge two adjacent 10-kb segments into one 20-kb segment. Based on the numbers of beads (*k_i_* and *k_i+1_*) of the two *i^th^* and (*i+1*)*^th^* merged parent 10-kb segments, we define the number of beads used to represent the resulted 20-kb segments, denoted as $\bar{k_{i\_20}}$ , according to the following:

$\bar{k_{i\_20}}=\frac{k_{i}+k_{i+1}}{2}$	(s10)

After the determination of bead numbers for all resulted 20-kb segments, we obtain the bead-on-a-string representation for the chromatin region of interest at 20-kb resolution. In the same manner, we generate the representation for the same region of 40-kb resolution from the representation of 20-kb resolution. We continue the procedure to generate the representations of lower resolutions, including 80-kb, 160-kb, 320-kb, 640-kb, 1280-kb, 2560-kb, and 5120-kb resolutions. After this procedure, we can determine the number of beads used to represent the chromatin region at lower resolutions based on the given (*k_1_*, *k_2_*,…*k_N_*).

(ii) Hierarchically generating 3D conformation of the chromatin region of interest at 10-kb resolution. We first generate an initial 3D structure of 5120-kb resolution of the chromatin region of interest by using random walk algorithm and then perform 200,000 steps of numerical integration to generate the simulated 5120-kb conformation.

Based on the coordinates of beads in the simulated 5120-kb conformation, initial coordinates of beads in the representation of 2560-kb resolution can be determined by using linear interpolation and are further relaxed with 200,000 steps of numerical integration. Upon subdivision, we also scale the bead coordinates at all three directions by 2 to keep so that the averaged radius of the beads (*b* = 1) is kept unchanged. $r_{s}$ and $E_{sphere}$ are also updated according to the change in *N* (but *b* is still kept at 1). In the same manner, the structure calculations are performed at 2560-kb, 1280-kb, 640-kb, 320-kb, 160-kb, 80-kb, 40-kb, 20-kb and finally 10-kb resolutions with the initial structure of each stage generated from the previous round of structure calculations. Such hierarchical simulation is performed for 50 times from 50 different initial 3D structures at 5120-kb resolution and finally generate 50 different simulated conformations of the chromatin region of interest at 10-kb resolution for a given vector (*k_1_*, *k_2_*,…*k_N_*). For each modeled chromatin region, we generated a group containing ten polymer models with each corresponding to a given vector (*k_1_*, *k_2_*,…*k_N_*). Therefore, we generate a total number of 500 conformations for the modeled chromatin region.

**Conformation ensembles generated in this paper**

The two conformation ensembles for K562 cell type predicted from different chromatin-accessibility data. These conformation ensembles cover the chromatin region including 22 autosomes and an X-chromosome. One ensemble is generated with the ATAC-seq data as input, and the other one with the DNase-seq data as input. Each ensemble contains 500 conformations of 10-kb resolution.

The conformation ensembles for IMR90 cell type. The conformation ensemble covers the chromatin region including 22 autosomes and an X-chromosome, is generated with the ATAC-seq data as input, and contains 500 conformations of 10-kb resolution.

The conformation ensembles for the cell types involved in the eight developmental stages of the differentiation from HSPCs to mature immune T cells. In these cell types, there are two chromatin regions of interest. One is the 10-Mb genomic region (Chr11:15Mb-25Mb) containing the regulator gene *Meis1* and the other is the 10-Mb genomic region (Chr12:105Mb-115Mb) enclosing the regulator gene *Bcl11b*. The eight cell types involved in the developmental stages of the differentiation are hematopoietic stem and progenitor cells (HSPCs), multipotent progenitor (MPP), common lymphoid progenitor (CLP), early T precursor (ETP), CD4 and CD8 double-negative 2 (DN2), DN3, DN4, and double-positive (DP) cells, respectively. For each region at each stage, we generate an ensemble containing 500 conformations at 10-kb resolution, with the DNase-seq experimental data as input.

The conformation ensemble (human cells) with chromatin fiber modeled as homopolymer. The conformation ensemble covers the chromatin region including 22 autosomes and an X-chromosome. Each chromosome is represented by polymer chain with homogenous DNA-packing density. Specifically, we set all the Poisson variables in (*k_1_*, *k_2_*,…*k_N_*) to be 0 and then each 10-kb segment is represented by one equal-volume bead.

**Determination of contact matrices** **from conformation ensembles**

Individual contact matrices of individual conformation. All the ensembles mentioned in this paper contain 500 individual 3D conformations of 10-kb resolution. According to the mentioned method of developing the polymer model of chromatin fiber, chromatin fibers of each conformation are divided into 10-kb segments with each 10-kb segment represented by *k* equal-volume beads. For each conformation, we measure the distance between the centroids of each pair of 10-kb segments. Based on the measurement, we build a contact matrix ***C*** for each conformation at the resolution of 10-kb, in which matrix element *c_ij_* is equal to 1 if the distance between the *i^th^* and *j^th^* segments is less than 4 bead diameters and *c_ij_* is zero when the distance is larger than 4 bead diameters.

Ensemble-average contact matrix. For each ensemble, we average the individual contact matrices across 500 conformations and obtain the ensemble-averaged contact matrix ***C* ^*^** which is defined as:

$c_{ij}^{*}=\frac{A_{ij}}{\sqrt{A_{i}*A_{j}}}$	(s11)

where $A_{ij}$ is the element of the matrix ***A*** which is generated by directly average individual contact matrices ***C*** across 500 conformations. Namely, the matrix ***A*** is defined as:

$A_{ij}=\frac{1}{500}\sum_{n=1}^{500} c_{ij,n}$	(s12)

Moreover, the $A_{i}$ and $A_{j}$ in the equation (s12) are calculated according to the formula below:

$A_{i}=\sum_{j} A_{ij}$ and $A_{j}=\sum_{i} A_{ij}$	(s13)

**Separation score**

Separation score at a 10-kb segment describes the degree of spatial segregation between chromatin on either side of the segment. In our study, we use separation score to identify domain boundaries in various contact matrices, including population-averaged Hi-C contact matrix, contact matrices of individual conformations, and ensemble-averaged contact matrix. Here, we use population-averaged Hi-C contact matrix ***C*** as an example to illustrate the calculation of separation score. First, we transform the population-averaged Hi-C contact matrix ***C*** into an arrowhead matrix ***M*** which is defined as:

$M_{i,i+d}=\frac{C_{i,i-d}-C_{i,i+d}}{C_{i,i-d}+C_{i,i+d}}$	(s14)

As shown in Figure S3, this transformation replaces domains with an arrowhead-shaped motif. Second, we define separation score based on the arrowhead matrix ***M***. As shown in Figure S3B we construct two edge-shared congruent right triangles, denoted as A and B, at each position along the diagonal line of the matrix ***M***. Separation score of the *i^th^* position, denoted as *S_i_*, is defined according to the following:

$S_{i}=\frac{1}{N_{A}}\sum_{i,j\in A} M_{ij}-\frac{1}{N_{B}}\sum_{i,j\in B} M_{ij}$	(s15)

where *N*_A_ and *N*_B_ are the total number of matrix elements in the triangles A and B, respectively. In our calculation, the length of horizontal edge of the two congruent right triangles is 250-kb and the length of the vertical edge (or shared edge) is 500-kb.

**Determination of the pairs of overlapping boundaries among Hi-C contact matrix and the simulated ensemble-average contact matrices**

To assess the performance of our model, we compare the genomic positions of TAD boundaries from experimental Hi-C contact matrix and positions of domain boundaries from the simulated ensemble-averaged contact matrix. We consider the two boundaries from different contact matrices as overlapping, if the distance between their genomic positions is less than 100 kb.

**Generation of random TAD boundaries**

The random domain boundaries were generated by circular shifting the genomic positions of domain boundaries determined from ensemble-averaged contact matrix along chromatin sequence.

**Determination of distance matrices** **from conformation ensembles**

Individual distance matrices of individual conformation. The calculation of the individual distance matrix at 30-kb resolution from individual conformations of 10-kb resolution is performed through two steps:

(i) Since each conformation of the conformation ensembles in this paper consist of consecutive 10-kb segments with each segment represented by *k* equal-volume beads, we sequentially merge three adjacent 10-kb segments into one 30-kb segments.

(ii) We measure the distance between the centroids of each pair of 30-kb segments and build an individual distance matrix for each conformation.

Ensemble-averaged distance matrix. For each ensemble, we directly average the distance matrices of individual conformations across 500 conformations and obtain the ensemble-averaged distance matrix.

**Calculation of distance matrix from fluorescence in situ hybridization experimental data**

Bintu *et al.* used multiplexed super-resolution fluorescence *in situ* hybridization (FISH) to image different chromatin regions, including K562 Chr21:29.38Mb-31.33Mb, IMR90 Chr21:29.38Mb-31.33Mb, IMR90 Chr21:20Mb-21.95Mb, HCT116(AUXIN treated) Chr21:36Mb-38.49Mb and HCT116(AUXIN untreated) Chr21:36Mb-38.49Mb. Based on the downloaded FISH data, we calculate the averaged distance matrix for each region over imaged cells. Since there are some nan values for coordinates of some chromatin segments in certain imaged cells, these coordinates are excluded in the averaged distance calculation.

**Identifying domain boundaries from the distance matrix derived from fluorescence *in situ* hybridization data**

We identify domain boundaries in the averaged distance matrix derived from FISH data by defining segregation value of each position. Specifically, the segregation value of each position is computed by averaging all the spatial distances between any pair of positions separately located in the two 240 kb regions on either side of the position. Repeating the calculation for each position, we can obtain the segregation values for all positions throughout the imaged region. The positions that are identified as boundaries should satisfy two criteria.: (i) the boundary position should show higher segregation value than any other positions within the 120 kb regions on either side of the position; (ii) the segregation value of boundary position should be higher than the averaged segregation value throughout the imaged region.

**Reference genome**

In this work, the reference genome hg19 is used for K562 and IMR90 cell line and the reference genome mm9 is used for cells of eight developmental stages differentiation from hematopoietic stem and progenitor cells (HSPCs) to mature immune T cells.
